# 2-Guanidino-quinazoline promotes the readthrough of nonsense mutations underlying human genetic diseases

**DOI:** 10.1101/2021.11.09.467859

**Authors:** Laure Bidou, Olivier Bugaud, Goulven Merer, Matthieu Coupet, Isabelle Hatin, Egor Chirkin, Pauline François, Jean-Christophe Cintrat, Olivier Namy

**Affiliations:** Université Paris-Saclay, CEA, CNRS, Institute for Integrative Biology of the Cell (I2BC), 91198, Gif-sur-Yvette, France; Sorbonne Université. CNRS; Université Paris-Saclay, CEA, INRAE, Département Médicaments et Technologies pour la Santé (DMTS), SCBM, 91191 Gif-sur-Yvette, France

**Author notes:** contributed equally (Laure Bidou, Olivier Bugaud).

## Abstract

Premature termination codons (PTCs) account for 10% to 20% of genetic diseases in humans. The gene inactivation resulting from PTC can be counteracted by the use of drugs stimulating PTC readthrough, thereby restoring production of the full-length protein. However, a greater chemical variety of readthrough inducers is required to broaden the medical applications of this therapeutic strategy. In this study, we developed a new reporter cell line and performed high-throughput screening (HTS) to identify potential new readthrough inducers. After three successive assays, we isolated 2-guanidino-quinazoline (TLN468). We assessed the clinical potential of this drug as a potent readthrough inducer on the 40 PTCs most frequently responsible for Duchenne muscular dystrophy. We found that TLN468 was more efficient than gentamicin, and acted on a broader range of sequences, without inducing the readthrough of natural stop codons.

## Introduction

The appearance of a nonsense mutation in a coding sequence creates a premature termination codon (PTC), preventing the correct production of the corresponding protein by interrupting translation and inducing the degradation of the transcript via the nonsense-mediated mRNA decay (NMD) pathway (Kim & Maquat, 2019). As a result, PTCs are associated with a large number of genetic diseases and cancers. A meta-analysis of nonsense mutations causing human genetic disease revealed that 11% of inherited diseases can be attributed to PTC mutations, 80% of which involve TGA and TAG (Mort *et al*, 2008).

In the last decade, considerable interest has focused on in-frame PTCs as potential therapeutic targets. Aminoglycosides (such as paromomycin, gentamicin, G418, and amikacin) and their derivatives (the NB series)(Leubitz *et al*, 2021; Sabbavarapu *et al*, 2016) have been shown to promote PTC readthrough by binding to mammalian ribosomes, leading to the partial restoration of full-length protein production in cultured mammalian cells and animal models (Amzal *et al*, 2021; Bidou *et al*, 2017; Prokhorova *et al*, 2017). The potential of this approach was first demonstrated *in vivo* by Barton-Davis and coworkers, who reported the restoration of dystrophin levels in the skeletal muscles of *mdx* mice to 10 to 20% those in wild-type animals, following subcutaneous injections of gentamicin (Barton-Davis *et al*, 1999). This strategy has been evaluated in a large number of genetic diseases (Lee & Dougherty, 2012), including cystic fibrosis (CF), and muscular dystrophies (Linde & Kerem, 2008), MPS I-H (Gunn *et al*, 2014), and several clinical trials have been report, with various degrees of success (Bordeira-Carrico *et al*, 2012; Keeling *et al*, 2014). Encouraging results have been obtained in some cases, particularly for mutations displaying high levels of readthrough in the presence of gentamicin (Sermet-Gaudelus *et al*, 2007). However, despite their medical value, aminoglycosides present adverse effects, with reports of various levels of ototoxicity and/or nephrotoxicity (Forge & Schacht, 2000; Mingeot-Leclercq & Tulkens, 1999). The development of novel, less toxic aminoglycoside analogs is considered a promising approach to overcoming this problem.

The limitations of aminoglycosides led to the development of other compounds with structures different from that of aminoglycosides. One such molecule, ataluren (also known as PTC124), was initially considered highly promising (Welch *et al*, 2007). Its clinical benefit remains a matter of debate, but it nevertheless recently obtained conditional approval from the EMA (Haas *et al*, 2015). Negamycin is a dipeptide that binds to the ribosomal A-site (Olivier *et al*, 2014), causing readthrough with a context dependence different from that of gentamicin (Pranke *et al*, 2018). Clitocine, a nucleoside analog, has been shown to have PTC readthrough activity (Benhabiles *et al*, 2017; Friesen *et al*, 2017), but its mechanism of action remains unknown. RTC/GJ compounds have been reported to effectively promote some readthrough activity with all three stop codons (Du *et al*, 2013), but this effect is not systematic (Gomez-Grau *et al*, 2015). Escin, a natural mixture of triterpenoid saponins isolated from horse chestnut (*Aesculus hippocastanum*) seeds, has recently been shown to promote readthrough of the G542X and W1282X mutations of the CFTR gene (Mutyam *et al*, 2016). Another molecule, 2,6-diaminopurine, has been shown to act as a specific corrector of UGA nonsense mutations, acting via the inhibition of Ftsj1, an enzyme that specifically modifies TRP-tRNA (Trzaska *et al*, 2020). Further studies will be required to determine the true clinical potential of this promising molecule.

The identification of new readthrough drugs is still of paramount importance, because most, if not all, the compounds identified to date display sequence specificity, restricting their potential clinical benefits to a limited number of patients. The identification of new drugs with different specificities will help to broaden the medical applications of this therapeutic strategy.

We set up a new experimental system for the high-throughput screening (HTS) of chemical libraries on mammalian cells without cell lysis, as a means of identifying new readthrough inducers. We performed two rounds of primary screening and then confirmed the readthrough activity of selected hits in a series of specific secondary assays for stop codon readthrough. The application of this approach to an initial panel of 17,680 initial molecules from two different chemical libraries led to the identification of one candidate model, which we named translectin (TLN468). We tested this molecule against the 40 different mutations of the DMD gene responsible for Duchenne myopathy. We showed that it stimulated PTC readthrough for a broad range of sequences. Finally, we used ribosome profiling to demonstrate that TLN468 does not induce readthrough on natural stop codons. This molecule displays no structural similarity to any known readthrough inducer, suggesting that it may be complementary to these other readthrough inducers or that it may promote readthrough for sequences not responding to other readthrough inducers.

## Results

### Development and validation of a stable cell line for readthrough inducer screening

We used an HTS approach to identify new readthrough compounds unrelated to known drugs. For the selection of drugs based on their readthrough activity, we first generated a stable mammalian cell line by integrating a secreted *Metridia* luciferase reporter gene interrupted by a nonsense mutation into NIH3T3 cells. This system combines the advantages of a live-cell assay with the sensitivity of an enzyme-based system. The coding sequence of the gene is interrupted by a nonsense mutation from *TP53*, R213X (TGA), embedded in its own nucleotide context. This mutation has the advantage of presenting an easily measurable basal level of readthrough (Floquet *et al*, 2011). We reasoned that if luciferase expression was actually due to readthrough of the R213X stop codon, then the addition of gentamicin should result in a significant increase in luciferase activity. We tested seven independent clones with a stable integration of the reporter gene in the presence and absence of 1.6 mM gentamicin. Significant induction was observed for all the clones (Figure S1). As expected, no induction of readthrough was observed in the presence of apramycin, an aminoglycoside known not to promote readthrough. We selected the clone displaying the strongest induction and the highest basal luciferase activity (C14) for further development (figure S1).

### Screening of two chemical libraries

During the development of the screening procedure, we found that coelenterazine (the substrate of the *Metridia* luciferase) was more stable at 18°C. We therefore performed the HTS at this temperature. We screened 17,680 molecules (16,480 from the Chembridge library and 1200 from the Prestwick library) for efficient stimulation of stop codon readthrough. We include eight negative controls (cells + DMSO) and eight positive controls (cells treated with 1.6 mM gentamicin). From this first screening, we selected 465 molecules that increased luciferase activity by a factor of at least 1.4 (figure 1.A), with a strictly standardized mean difference (SSMD) of at least 2 (Zhang *et al*, 2007). The SSMD is used in HTS as a means of identifying compounds with a defined magnitude of difference from the median values obtained in the assay; values above 2 are considered to indicate a strong effect (Zhang, 2011). These first hits were then subjected to a second round of screening in the same conditions, with duplicates for each molecule. We retained 43 molecules based on their ability to induce luciferase activity by a factor of at least 2 relatives to untreated cells.

**Figure 1:**
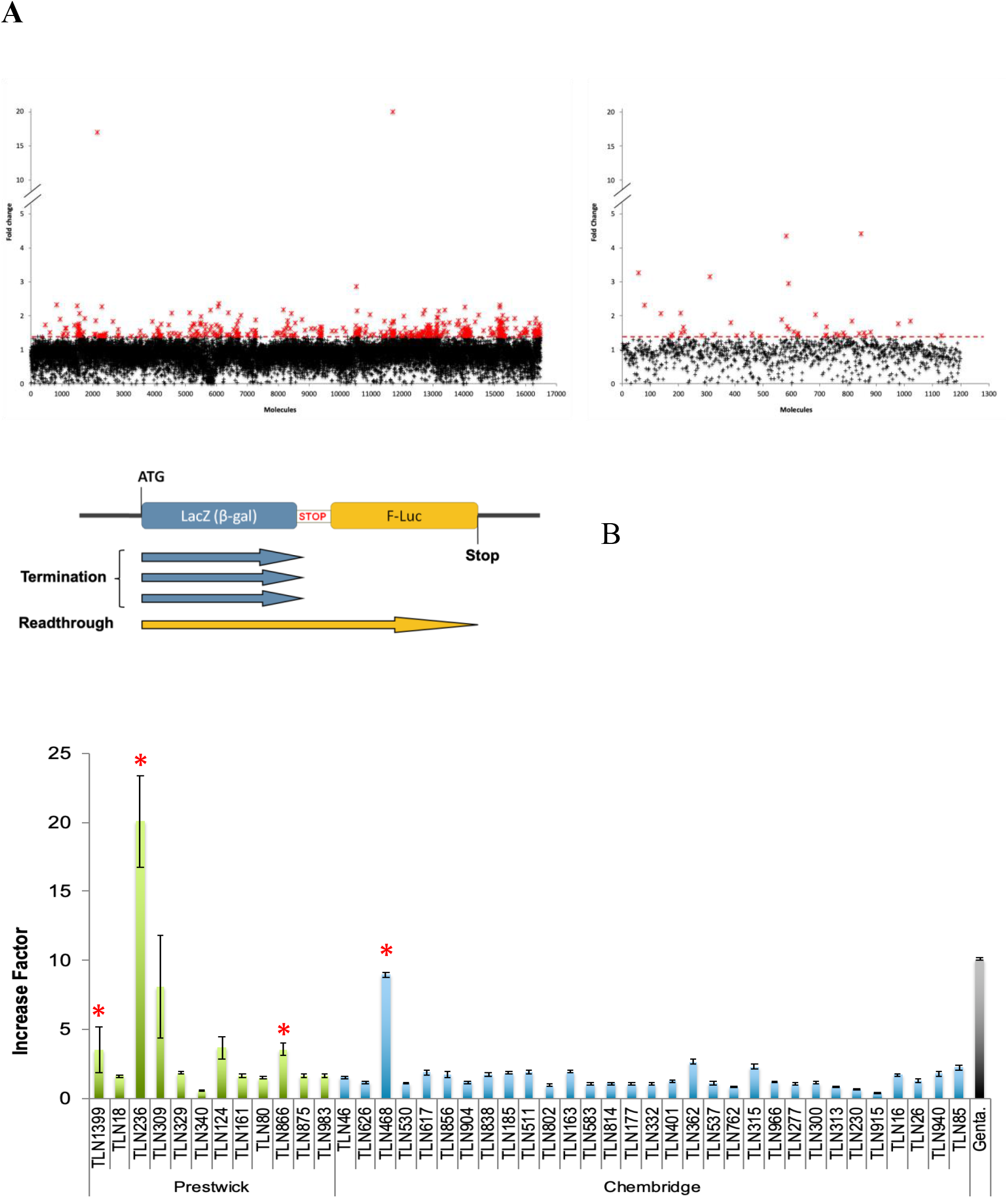
Primary selection and secondary quantification of the readthrough effect of the positive hits. **A)** Positive hits were selected according to the fold-change in luciferase activity between treatment (at 50 μM) and the median value for the negative controls (untreated cells). The molecules from the Chembridge chemotherapy library and those of the Prestwick library are represented in the left and right panels, respectively. The red crosses correspond to the 465 molecules inducing at least a 1.4-fold increase in activity. **B)** The dual reporter system is illustrated in the top panel. The red stop indicates the R213X sequence inserted between *lacZ* and *F-luc*. The ability of the 43 retained molecules to stimulate readthrough is illustrated in the lower panel. NIH3T3 cells were treated for 24 hours with gentamicin (2.5 mM) as a control or with one of the 43 molecules selected (50 μM) from the Prestwick chemical library (green bars) or the Chembridge chemical library (blue bars). The red asterisks represent the 4 molecules selected for further characterization.

Such screening can lead to the identification of many false-positive hits. We subjected the 43 molecules to several independent assays, to limit the selection of false positives and ensure the selection exclusively of *bona fide* readthrough inducers with a clear effect on stop codon readthrough.

### Quantification of stop codon readthrough

The initial screening method was highly sensitive and very convenient, but subject to several inherent biases. Indeed, any molecule increasing the production (mRNA transcription, stability, translation) or secretion of the Met-luciferase will lead to the identification of a false-positive hit. We got round this problem by assessing the ability of each drug to stimulate readthrough in a dual reporter system, to quantify stop codon readthrough efficiency (Figure 1.B)(Bidou *et al*, 2004). This reporter system includes enzymatic activities (β-galactosidase and firefly luciferase) different from that used in the initial screen. Furthermore, β-galactosidase is used as an internal control for the normalization of expression levels. The use of this second reporter system therefore made it possible to eliminate many of the false-positive hits selected during the initial screen. For each molecule tested, three independent measurements were performed (Figure 1.B). Six molecules induced at least a three-fold increase in PTC readthrough (Figure 1.B). Only four of these molecules were available at a larger scale. This second reporter system eliminated false-positive hits very efficiently, but it was nevertheless a reporter system based on enzymatic activity. We decided to use a more physiological system to evaluate the last four potential hits.

### Restoration of TP53 protein production in HDQP-1 cells

We used the human HDQ-P1 cell line, which carries the R213X nonsense mutation in its endogenous TP53 gene (Wang *et al.*, 2000). We assessed production of the full-length p53 by western blotting. The advantages of this third system are that it makes use of an endogenous nonsense mutation and direct visualization of the final product induced by the potential hit (i.e. the full-length protein). Two molecules (TLN468 and TLN309) clearly stimulated the production of full-length p53. Interestingly, we simultaneously observed an accumulation of the truncated form potentially corresponding to a stabilization of the TP53 mRNA due to the inhibition of NMD. For confirmation of this observation, we performed RT-qPCR on the TP53-R213X mRNA. Our results showed that this mRNA was stabilized five-fold by TLN468 but only two-fold by TLN309 (Figure 2.B).

**Figure 2:**
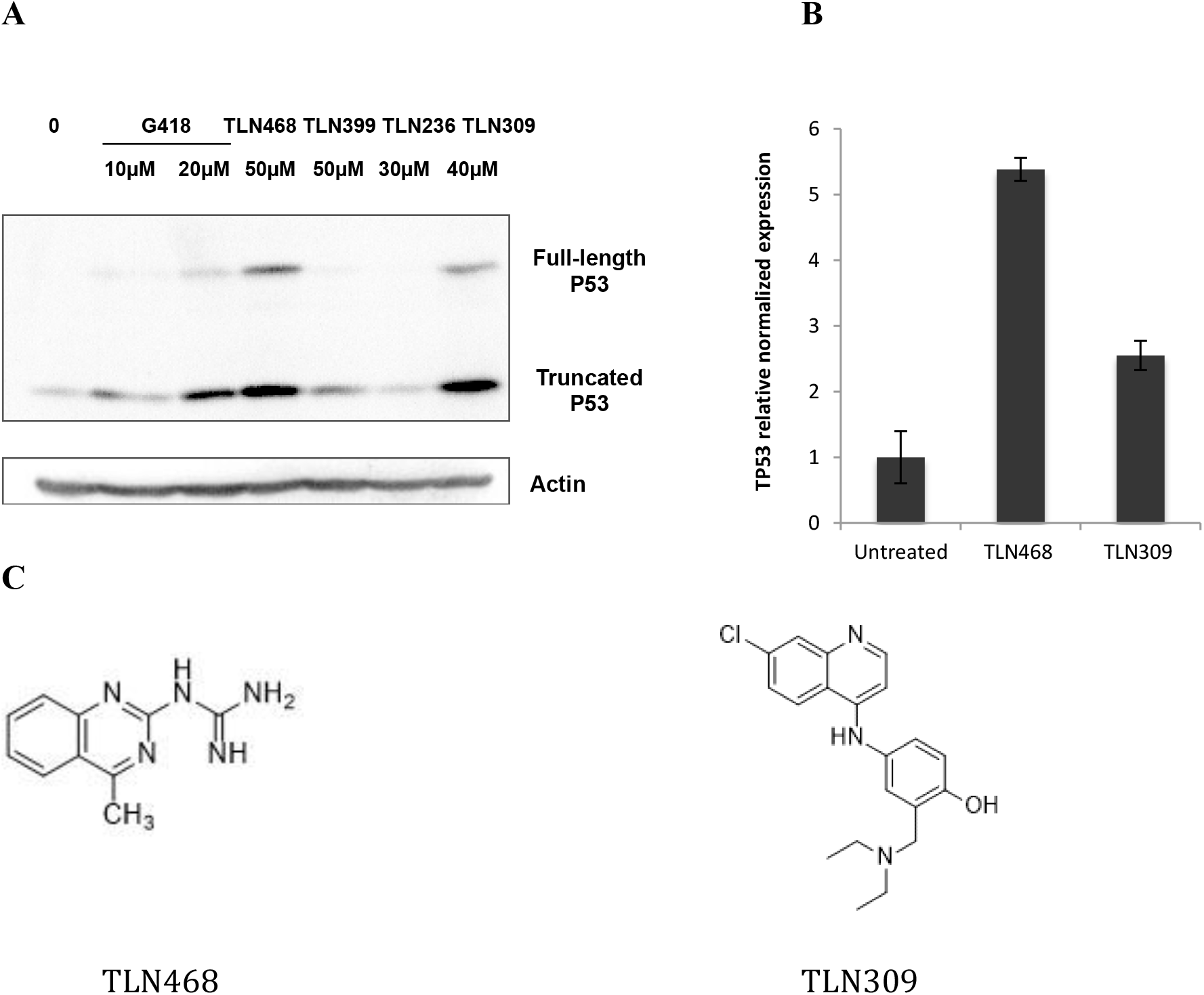
Effects of the four selected hits on the endogenous nonsense R213X mutation of *p53*. **A)** Detection by western blotting of the full-length p53 protein in HDQ-P1 cells carrying the endogenous nonsense mutation R213X. Cells were left untreated (0) or were treated with G418 or the four positive hits for 48 h. Membranes were probed with the DO-1 antibody directed against the N-terminus of p53, and an anti-actin antibody was used as a loading control. **B)** Mutant p53 mRNA levels were determined by quantitative PCR (*n* = 3). HDQ-P1 cells were left untreated or were treated with TLN468 (80 μM) or TLN309 (40 μM) for 48 hours. The results are expressed relative to the normalized relative amount of mRNA in the absence of a treatment. **C)** Structures of TLN468 and TLN309 (amodiaquine).

A structural review of these two last candidates (Figure 2.C) indicated that TLN309 was a 4-aminoquinoline (amodiaquine) similar to chloroquine, with autophagy-lysosomal inhibitory activity, promoting a ribosome biogenesis stress (Espinoza *et al*, 2020; Qiao *et al*, 2013). We therefore decided to pursue our analysis exclusively with TLN468, a 2-guanidino-quinazoline that we named translectin (Figure 2.C).

### Restoration of functional TP53 expression by stop codon readthrough

TLN468 was identified as a very encouraging hit for restoring protein production from a gene interrupted by a nonsense mutation. The production of a full-length protein is essential, but not sufficient to validate the correction of the defective gene and the functionality of the readthrough protein. Indeed, we have shown that at least three tRNAs (Tyr, Gln, Lys) can be used to read through the UAG codon (Blanchet *et al*, 2018). The precise amino-acids incorporated may considerably modify the activity of the restored full-length protein. We therefore used two different systems to determine whether the full-length p53 produced in the presence of TLN468 was functional. As p53 is a transcriptional activator, we first used a plasmid carrying a luciferase gene under the control of a p53 promoter (Figure 3.A). We then used RT-qPCR to quantify the Bax mRNA, one of the major cellular targets of p53 (Figure 3.B). TLN468 treatment induced a dose-dependent increase in p53-dependent luciferase activity, consistent with the 2.5-fold increase in Bax mRNA levels in the presence of 80 μM TLN468. These findings confirm at least a partial restoration of the p53 activity. Overall, these independent assays indicate that TLN468 is a promising readthrough inducer.

**Figure 3.**
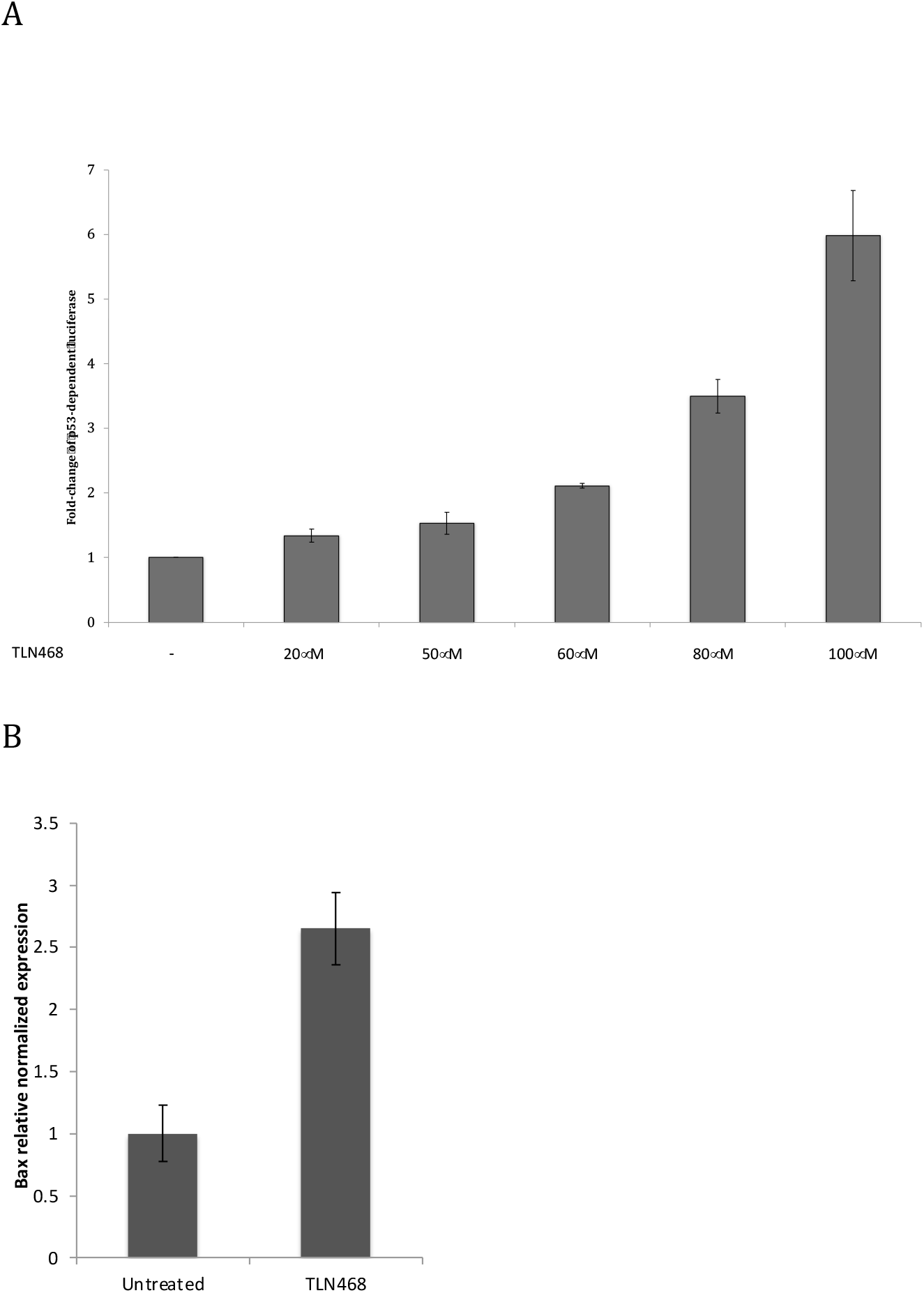
TLN468 restores p53-R231X activity. **A)** Human H1299 (p53^−/−^) cells were cotransfected with pCMV-R213X, pCMV-LacZ and p53BS-luc. The pCMV-R213X vector expresses the P53 gene with the nonsense R213X mutation. The p53BS-luc vector expresses the luciferase gene under the control of a p53-dependent promoter, leading to the detection of a transcriptionally active p53. The pCMVLacZ vector was used for normalization. **B)** HDQ-P1 cells were left untreated or were treated with TLN468 (80 μM) for 48 hours. mRNA levels for Bax, one of the major targets of p53, were determined by reverse transcription-quantitative PCR (*n* = 3). The results are expressed relative to the normalized relative amount of mRNA in the absence of treatment, set to 1.

We used the same mutation (R213X) in all these assays. Having established that TLN468 induced stop codon readthrough for this particular premature stop codon, we then investigated its spectrum of action against a range of nonsense mutations.

### Specificity of TLN468

We addressed the question of TLN468 specificity using the 40 most frequent premature nonsense mutations found in DMD gene and responsible for Duchenne and Becker muscular dystrophy (Table S1; Figure 4.A). Each nonsense mutation (+ 9 nucleotides in each side) was inserted into the dual reporter construct described above. Stop codon readthrough efficiency was quantified with or without gentamicin or TLN468, in six independent experiments. TLN468 promoted at least a doubling of stop codon readthrough for 36 of the 40 sequences tested (Figure 4.B). Interestingly, it also outperformed gentamicin for 90% of these sequences. We conclude that TLN468 is active against a wide variety of sequences, but that its action is sequence-dependent, as for most other known readthrough inducers. This sequence specificity seems to be different from that of gentamicin, for which a C in the +1 position relative to the PTC is preferred (Figure 4.C). This observation led us to investigate whether any additive or synergistic effects occurred when gentamicin and TLN468 were added together. We performed the assay with the dual reporter system carrying the R213X mutation and two concentrations of gentamicin. The two drugs clearly had an additive effect on stop codon readthrough (Figure 5).

**Figure 4:**
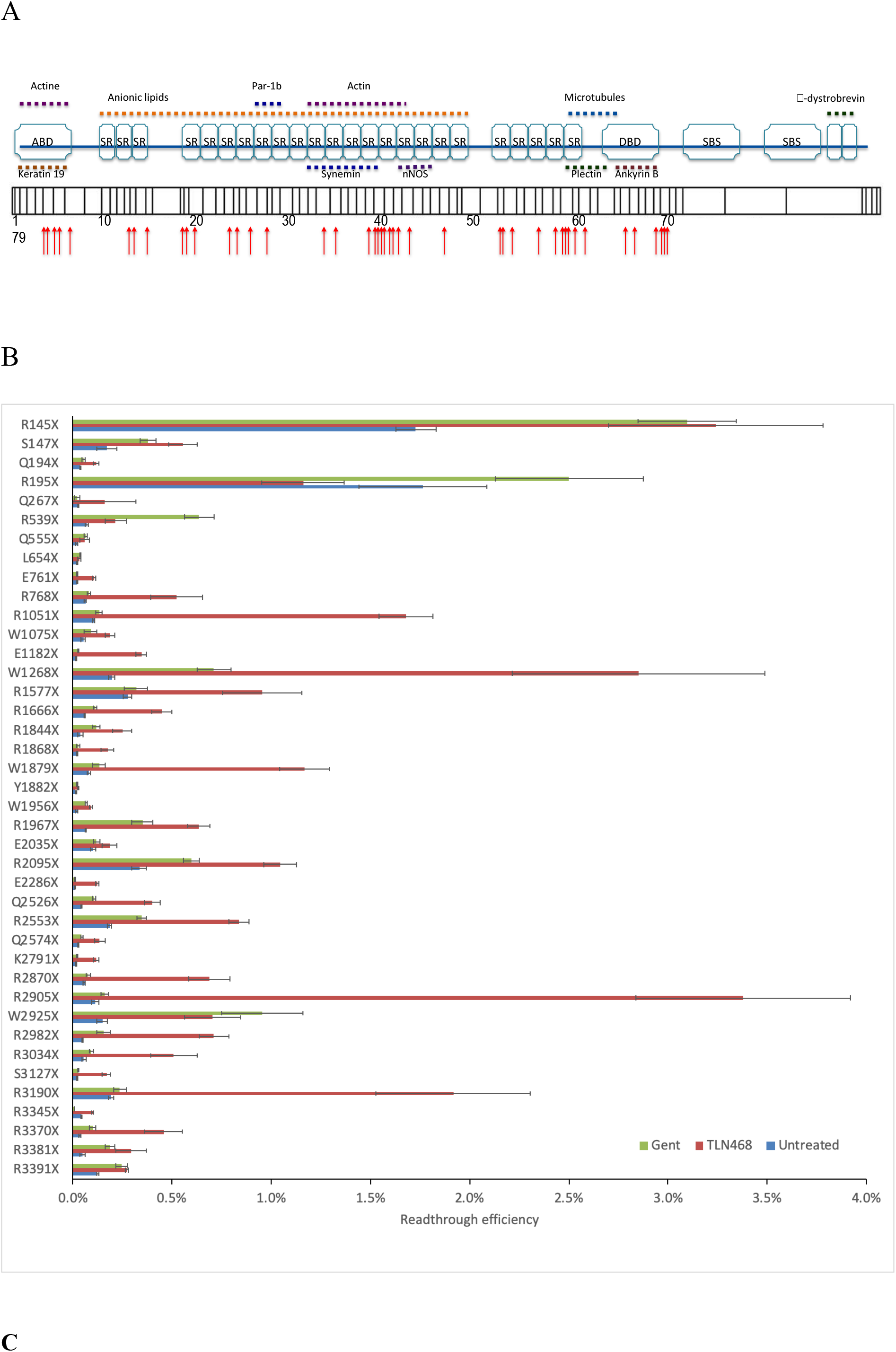
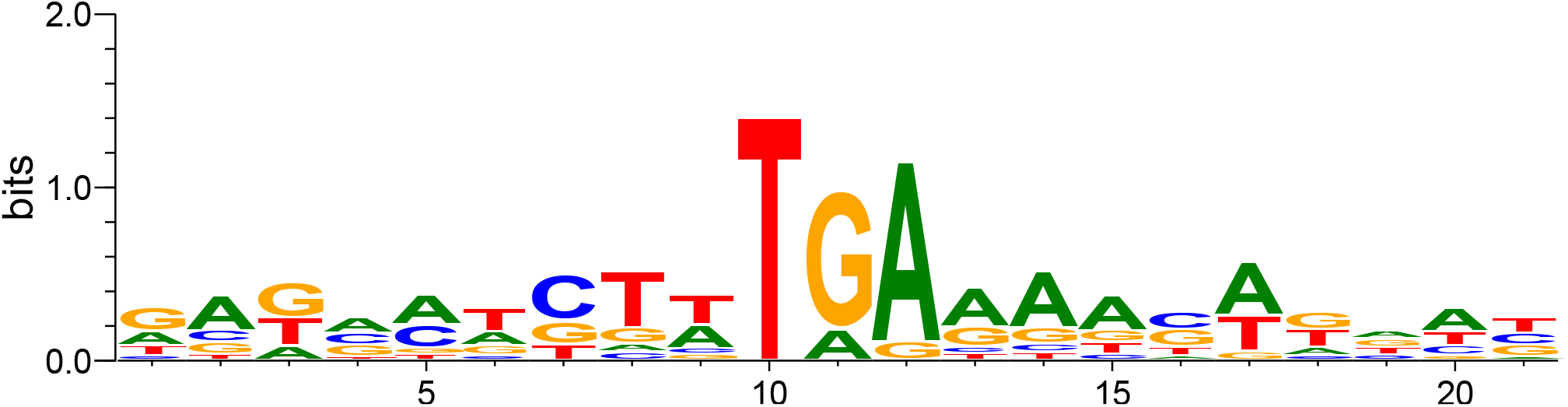
Sequence specificity of TLN468. A) Schematic representation of the dystrophin gene and its different domains with the position of the selected 40 mutations. B) Quantification of stop codon readthrough efficiency for the 40 most frequent nonsense mutations found in the DMD gene (*n*=6), in the presence or absence of 80 μM TLN468. A similar quantification was performed with gentamicin (2.5 mM), in the same conditions, to compare the effects of TLN468 with those of gentamicin. C) Logo obtained from the 11 sequences displaying the strongest TLN468 induction (generated with WebLogo 3).

**Figure 5:**
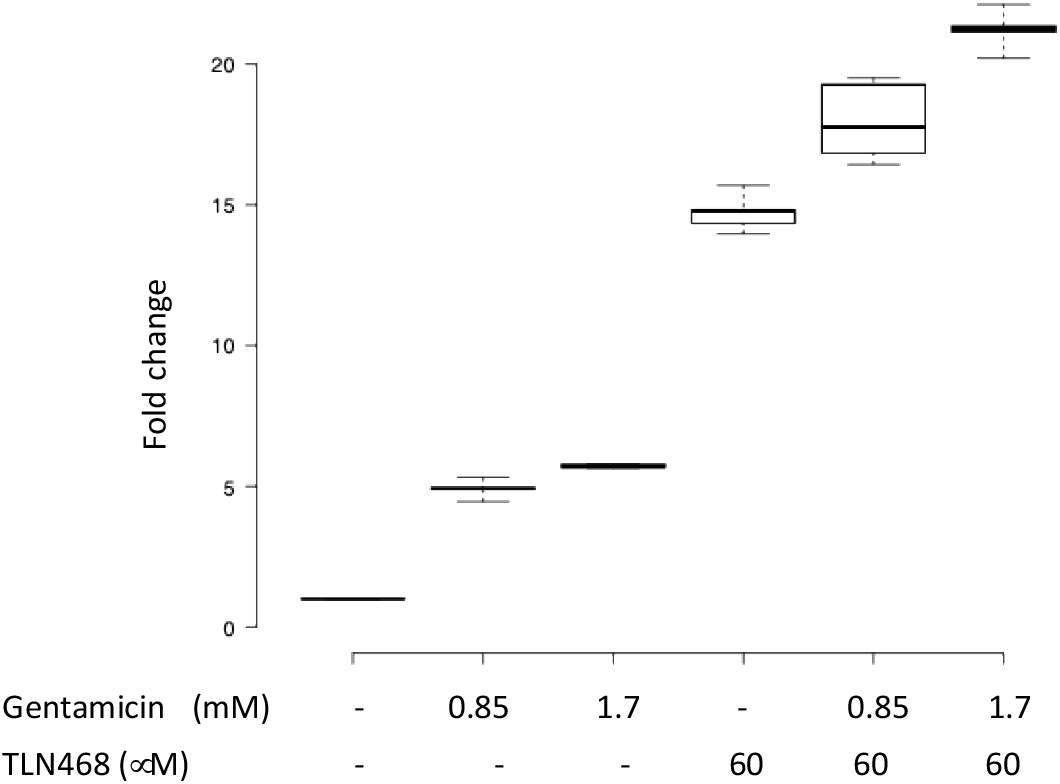
Additive effect of TLN468 and gentamicin. HeLa cells were treated with gentamicin, TLN468 or both drugs for 24 h. Stop codon readthrough (R213X) was then quantified with the dual reporter system. The values for the untreated cells are set to 1 and fold-changes are calculated relative to this value. At least six independent measurements were performed for each treatment.

### Genome-wide analysis of the action of TLN468

We assessed the genome-wide effect of TLN468 on translation termination, by performing ribosome profiling in cells treated with this compound. HeLa cells were treated for 24 h with TLN468 (80 μM), DMSO as a negative control or G418 (gentamicin) as a positive control. We checked the quality of the data for periodicity and coding sequence (CDS) enrichment, and we then checked for global readthrough at normal stop codons by performing a metagene analysis, aligning all transcripts from their stop codons backwards (Figure 6A). Consistent with published data, we detected a strong accumulation of ribosome footprints at the stop codons (Wangen & Green, 2020). Interestingly, G418 treatment led to the emergence of a periodic signal immediately downstream from the stop codon, whereas no signal was observed in the absence of this drug. This reflects the induction of readthrough at natural stop codons by G418, as previously described (Wangen & Green, 2020). By contrast, TLN648 did not promote genome-wide stop-codon readthrough.

**Figure 6:**
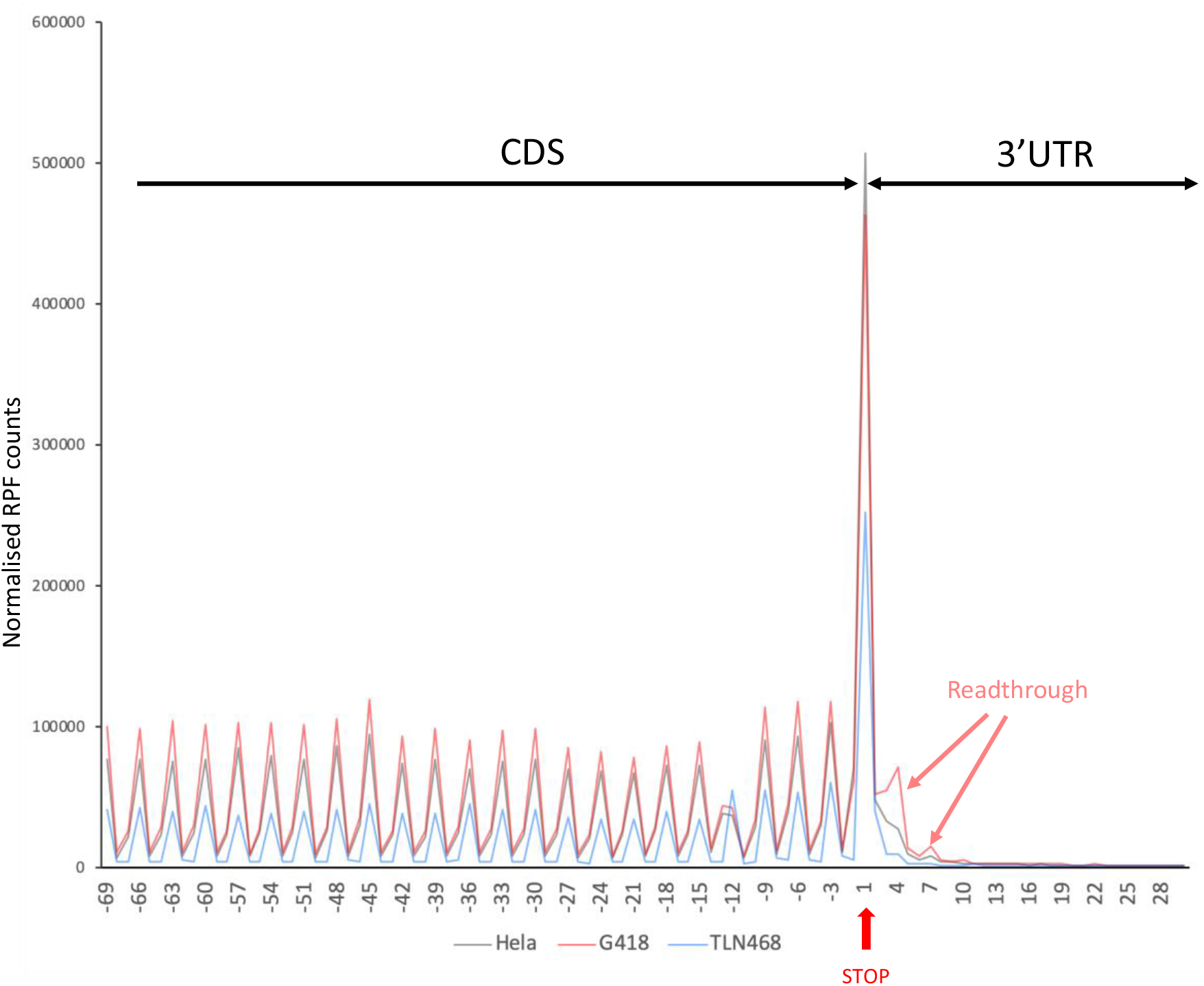
Metagene analysis of the action of TLN468 based on ribosome profiling. Ribosome profiling was performed in HeLa cells treated for 24 h with TLN468 (80 μm) (blue), DMSO (gray) as a negative control or G418 (red)(722 μM) as a positive control. Each individual transcript was aligned from the stop codon back, and RPF are indicated by the first nucleotide at the A-site. The vertical red arrow indicates the first position of the stop codon numbered +1, +2, +3. RPF corresponding to ribosomes reading through natural stop codons are also indicated.

## Discussion

We screened 17,680 compounds to identify drugs promoting nonsense suppression. We used two different chemical libraries: the Prestwick library, which contains drugs already approved by the FDA and the EMA, and a subset of the Chembridge small-molecule screening library, which contains drug-like and lead-like screening compounds with a wide range of chemical diversity.

We used a three-step workflow strategy based on complementary assays collectively assessing the readthrough specificity of the compounds. For the initial screening, we constructed a completely new reporter cell line expressing a secreted luciferase encoded by the *Metridia* luciferase gene interrupted by the R213X mutation (Floquet **et al.**, 2011). Absolute values for luciferase activity can be impacted at various steps of its expression may be affected by the various steps in the production and action of the enzyme (transcription, translation, RNA & protein stabilities), but the secretion of the enzyme in this assay greatly simplifies the screening procedure by overcoming the need for a cell lysis step. After two rounds of screening with this system, we selected 43 compounds yielding a robust and reproducible induction of luciferase activity.

In our secondary screen, we used a dual-reporter system (Bidou **et al.**, 2004) for the accurate quantification of stop codon readthrough. This second reporter system eliminated a large number of false positives, because it was based on different enzyme activities (not all compounds affecting the *Metridia* luciferase or its substrate would be able to affect this second reporter), and because it allowed internal normalization against lacZ to eliminate variability in mRNA abundance or translation initiation. After this second screen, we selected four compounds for further investigation (Figure 1.B).

For the third screen, we decided to test the compounds in the human HDQ-P1 cell line carrying an endogenous PTC mutation in the *TP53* gene (Wang **et al**, 2000). This system provided us with an opportunity to assess the effect of the drugs in a situation in which the PTC was present in a genetic environment different from that in the reporter systems. On western blots, we clearly detected the short p53 isoform resulting from the premature arrest of translation at the PTC, and the full-length p53 following the induction of PTC readthrough by the drugs (Figure 2.A). In parallel with the appearance of the full-length protein, we also observed an increase in the amount of the truncated form of p53. This was expected, as PTC readthrough inhibits NMD, the process that degrades PTC-containing mRNAs (Allamand *et al*, 2008; Bidou **et al.**, 2017; Keeling *et al*, 2004). The stabilization of the TP53 mRNA in the presence of TLN468 and, to a lesser extent, in the presence of TLN309, was further confirmed by RT-qPCR (Figure 2.B). TLN309 is known as amodiaquine, an antimalarial drug that stabilizes p53 via the RPL5/RPL11-5S rRNA checkpoint (Espinoza **et al.**, 2020). This may also account for the detection of the full-length p53 in the presence of this drug in our assay. After these three successive screens, we selected a single candidate: 2-guanidino quinazoline (TLN468)(Figure 2.C).

We focused on TLN468, to determine its therapeutic potential. We first showed that treating human cells carrying a defective p53 with various amounts of TLN468 led to the restoration of a functional p53 (Figure 3) and activation of the expression of BAX, one of the major targets of p53 (Riley *et al*, 2008). This demonstrates that TLN468 can restore the biological function of a gene carrying a PTC when added to cell culture. One key determination is that of the specificity of action of TLN468 according to the PTC tested and the surrounding nucleotides. We addressed this question by performing a systematic readthrough analysis on the 40 most frequent nonsense mutations of the DMD gene underlying Duchenne muscular dystrophy in the presence of either gentamicin or TLN468 (Figure 4). As previously reported for other PTCs, basal readthrough levels are highly variable, ranging from 0.01% for Y1882X and E2286X to 2% for R145X and R195X. Interestingly, R145X (<u>UGA</u>CAAU) and R195X (<u>UGA</u>CUGG) are found in a context very similar to the most efficient readthrough context identified in human cells (UGACUAG) (Loughran *et al*, 2014), whereas Y1882X (<u>UAA</u>AAGA) and E2286X (<u>UAA</u>GAGU) present no similarity to this efficient readthrough motif. As previously reported (Floquet *et al*, 2012), we found no correlation between the basal readthrough level and the level of induction by either gentamicin or TLN468. We observed a clear sequence-specificity for TLN468, which differed from that of gentamicin. By analyzing the 11 sequences leading to the strongest induction of TLN468, we found that an A at position +1 relative to the stop codon was the most favorable context for TLN468 action (Figure 4C), whereas a C in the +1 position was the context resulting in the highest readthrough efficiency in the presence of gentamicin (Floquet **et al.**, 2012; Manuvakhova *et al*, 2000). However, we cannot exclude a sequence bias due to the relatively small number of sequences analyzed. The atlas we generated for gentamicin and TLN468 will also be of considerable interest for medical applications of gentamicin, as it will indicate which mutations are likely to respond to gentamicin treatment for translational stop codon suppression. Finally, we investigated the possibility of using TLN468 and gentamicin together to increase the therapeutic benefit. We tested the addition of two concentrations of gentamicin to HeLa cells with and without TLN468 (Figure 5). We observed a clear additive effect of the two drugs, suggesting that this combination should be considered as a means of improving overall treatment efficacy. This complementarity of action is very interesting medically, but it also indicates that the two compounds do not bind to the same site. We cannot exclude the possibility that TLN468 binds elsewhere on the ribosome, particularly given that 2-guanidino-quinazoline has been described as a bacterial translation inhibitor with antibacterial activity (Komarova Andreyanova *et al*, 2017). These characteristics are highly reminiscent of those of gentamicin, which is used primarily as an antibiotic. Genome-wide RiboSeq analysis clearly indicated that TLN468 did not induce stop codon readthrough at natural stop codons. We cannot exclude the possibility of a weak impact on some specific genes, but this finding is nevertheless consistent with the absence of a global deregulation of translation termination. Overall, our data suggest that TLN468 acts through a mechanism different from that of aminoglycosides, resulting in the specific stimulation of PTC readthrough with no alteration of normal termination process.

## Materials and Methods

### Cell lines and plasmids

All cells were cultured in DMEM plus GlutaMAX (Invitrogen), except for H1299 cells, which were cultured in RPMI plus GlutaMAX (Invitrogen). The medium was supplemented with 10% fetal calf serum (FCS, Invitrogen) and 100 U/mL penicillin/streptomycin. Cells were incubated in a humidified atmosphere containing 5.5% CO_2_, at 37°C. NIH3T3 cells are embryonic mouse fibroblasts. H1299 is a p53-null cell line established from a human lung carcinoma (provided by the ATCC). HDQ-P1 is homozygous for a nonsense mutation at codon 213 (CGA to TGA) of the p53 gene. This cell line was established from a human primary breast carcinoma and was provided by the DSMZ-German Collection of Microorganisms and Cell Cultures. For the generation of a stable mammalian cell line, we used a secreted *Metridia* luciferase reporter gene derived from pMetLuc2 (Clontech). The coding sequence of this gene was interrupted by the TP53 nonsense mutation R213X, located in its own nucleotide context and inserted into an *Eco*53KI site created by directed mutagenesis at nucleotide 57. The final construction was named pML213 and was used for the stable transfection of NIH3T3 cells with the JetPei reagent (Invitrogen). We tested the capacity of seven neomycin-resistant clones to express active *Metridia* luciferase after readthrough induction in the presence of 1.6 mM gentamicin for 24 hours. We then removed 50 μL of culture medium from each well and incubated it in the presence of substrate: coelenterazine, according to the conditions recommended by the supplier (Ready-To-Glow Secreted Luciferase Reporter Assay; Clonetech). The photon emission generated by the reaction was measured in a plate luminometer (Tecan).

The clone presenting the highest increase factor between the treated and control conditions was selected for the HTS.

### HTS screening

3T3pML213 (C14) cells were cultured in 60 cm^2^ cell-treated culture dishes in DMEM Glutamax (Life Techologies) medium (10 mL) supplemented with serum (10%). The cells were incubated at 37°C, under an atmosphere containing 5% CO_2_. The cells were detached by trypsin treatment (2 mL/plate) for five minutes at 37°C under an atmosphere containing 5% CO_2_. The cells were then transferred to DMEM supplemented with 10% serum in an Erlenmeyer flask, and were incubated with slow rotation. The density of the cell suspension was determined by five cell counts with a Scepter (Millipore) and 60 μm probes. The density of the suspension was adjusted to 25,000 cells per mL. The cells were then automatically plated in 96-well plates (Sterile Corning® 96-well flat clear bottom white polystyrene TC-treated microplates) at a seeding density of 2500 cells/well. Initial screening was conducted on a drug repurposing library, the Prestwick Chemical Library® (1,280 small compounds already approved by the FDA, EMA and other agencies). We then extended the screening to a subset of the Chembridge DIVERset® library comprising 16,480 compounds. Each 96-well plate contained 80 different compounds (initial concentration, 10 mM in DMSO) and 16 controls (columns 1 and 12). Gentamicin (2 mg/mL) was used as a positive control and the negative control was 0.5% DMSO. Each molecule from the two libraries was tested at a concentration of 50 μM, with overnight incubation. Screening and secondary validations of both the Prestwick® and Chembridge DIVERset® libraries were performed according to the same procedure (see below).

In a dark room, at a temperature below 16°C, coelenterazine (Nanolight) was dissolved at a concentration of 10 mM in degassed (N_2_) ethanol. A substrate/buffer mixture was then prepared at a concentration of 125 μM in a 15 mL tube to limit the area available for oxidation. This mixture was degassed by bubbling with dry nitrogen and was incubated for five minutes at 4°C. We then dispensed 2 mL of the mixture into the first row of a 6 mL 48-well plate. A multichannel pipetted was used to transfer 280 μL of the mixture to a conical-bottomed plate, to minimize the dead volume. This step was immediately followed by centrifugation at 500 x *g* for 30 seconds. We then deposited 70 μL of mineral oil (Sigma) on the surface of the mixture in the substrate plate, using a 96-channel Liquidator dispenser. The plate was incubated at 4°C. Using the Liquidator, we removed 10 μL of the mixture and deposited it in the screening plate. The contents of the plate were immediately manually homogenized with an orbital motion and the plate was placed in a Spectramax M5e plate luminometer (Molecular Device) for reading. All hits from the initial screening were ordered in powder form and retested.

### Statistical validation of the new reporter cell line

For validation of our screening strategy, we first applied the screening protocol to five 96-well plates, using gentamicin as the positive control and DMSO as the negative control. For statistical validation, we used two parameters described by Zhang **et al.** in 1999. The Z factor provides an easy and useful summary of assay quality and is a widely accepted standard (Zhang *et al*, 1999). The Z factor combines information about both the location and scale of the distributions of the sample signal and background. A Z factor greater than 0.5 is often interpreted as an indication of acceptable assay quality. Another powerful parameter proposed by Zhang is the strictly standardized mean difference (*SSMD*), which is robust to both measurement unit and strength of the positive control. It takes into account data variability in the two groups and integrates a probability interpretation. We retained drugs obtaining an SSMD score ≥ 2 during the screening procedure and a Z factor of at least 0.5 during the validation of our screening protocol.

### Readthrough quantification

For each nonsense mutation tested, complementary oligonucleotides corresponding to the stop codon and nine nucleotides on either side of the stop codon were ligated into the pAC99 dual reporter plasmid, as previously described (Bidou **et al.**, 2004). This dual reporter plasmid was used to quantify stop codon readthrough through the measurement of luciferase and beta-galactosidase (internal normalization) activities, as previously described (Stahl *et al*, 1995). Readthrough levels for nonsense mutations were analyzed in the presence or absence of the tested molecules. The cells were used to seed a six-well plate. The following day, they were transfected with the reporter plasmid in the presence of JetPei reagent (Invitrogen). They were incubated for 14 h and then rinsed with fresh medium, with or without potential readthrough inducers. Cells were harvested 24 hours later, with trypsin–EDTA (Invitrogen), lysed with Glo lysis buffer (Promega) and beta-galactosidase and luciferase activities were assayed as previously described (Stahl **et al.**, 1995). Readthrough efficiency was estimated by calculating the ratio of luciferase activity to beta-galactosidase activity obtained with the test construct, with normalization against an in-frame control construct. At least five independent transfection experiments were performed for each assay.

### RNA extraction and RT-qPCR

For the analysis of mRNA levels for *p53* and its transcriptional target gene, *Bax*, we extracted total RNA from HDQ-P1 cells that had or had not been treated with G418 (400 μM) or TLN468 (80 μM) for 48 hours (RNeasy Mini Kit, Qiagen). The RNA was treated with DNAse I (RNase-free DNase) and quantified with a Denovix ds-11 spectrometer. The absence of RNA degradation was confirmed by agarose gel electrophoresis. The first-strand cDNA was synthesized from 2 μg of total RNA, with random primers and the SuperScript II Reverse Transcriptase (Invitrogen), as recommended by the manufacturer. Quantitative PCR was then performed with equal amounts of the various cDNAs, with a CFX96 thermocycler (Biorad), and the accumulation of products was monitored with the intercalating dye FastStart Universal SYBRGreen Master (ROX) (Roche). We quantified mRNA levels relative to three reference mRNAs: RPL32, Hprt1 and HMBS. In each experiment, results are expressed relative to those for untreated cells, for which the value obtained was set to 1. Relative levels of gene expression were calculated at early stages of PCR, when the amplification was exponential and might, therefore, be correlated with the initial number of copies of the transcript. The specificity of quantitative PCR was checked by agarose gel electrophoresis, which showed that a single product of the desired length was produced for each gene. A melting curve analysis was also performed. Single product-specific melting temperatures were identified for each gene. For the quantification of each mRNA, three independent experiments (from biological replicates) were performed in triplicate. We used the following oligonucleotide pairs for amplification: p53 forward: 5’CCGCAGT CAGATCCTAGCG 3’ and reverse: 5’CCATTGCTTGGGACGGCAAGG 3’ ; Bax forward : 5’GCTGTTG GGCTGGATCCAAG 3’ and reverse 5’ TCAGCCCATCTTCTTCCAGA

### Western-blot analysis

HDQ-P1 cells (R213X) were treated with G418 (10 and 20 μM), or the selected molecules (50 μM TLN468, 50 μM TLN399, 30 μM TLN236 and 40 μM TLN309), for 48 h. Cells were harvested by treatment with trypsin–EDTA (Invitrogen), lysed in 350 mM NaCl, 50 mM Tris– HCl pH 7.5, 1% NP-40, and protease inhibitor cocktail (Roche) and disrupted by passage through a syringe. Total proteins were quantified with Bradford reagent (Biorad) and extracts were denatured by incubation in Laemmli buffer for 5 minutes at 90°C. We subjected 30 μg of total protein from HDQ-1 cells to SDS–PAGE in 4/10% Bis–Tris gels. Proteins were transferred onto nitrocellulose membranes, according to the manufacturer’s instructions (Biorad). Membranes were saturated by overnight incubation in 5% skimmed milk powder in PBS, and incubated for 1 hour with the primary monoclonal antibody, DO-1 (N-terminal epitope mapping between amino-acid residues 11 and 25 of p53; Santa Cruz Biotechnologies, 1/400) or a monoclonal antibody against mouse actin (Millipore, 1/2000). After three washes in PBS supplemented with 0.1% Tween, the membranes were incubated with the secondary antibody [horseradish peroxidase-conjugated anti-mouse IgG (1/ 2500) for 45 minutes. The membranes were washed five times and chemiluminescence was detected with ECL Prime Western Blotting Detection Reagents (Amersham, GE Healthcare). The signal was quantified with ImageJ software.

### TP53 protein activity assays

We investigated the transcriptional activity of the p53 protein in H1299, a p53-null cell line. Cells were cotransfected, by the JetPei method, with the p53BS-luc reporter plasmid containing the firefly luciferase gene downstream from seven p53 binding sites, pCMVLacZ and pCMVp53R213X containing the p53 cDNA interrupted by the R213X stop codon. TLN468 (20, 50, 60, 80 and 100 μM) was added to the medium just before transfection, for a total of 20 hours of treatment. Protein extracts were then prepared and enzymatic activities were measured. Transfection with pCMVLacZ was used to normalize transfection efficiency, cell viability and protein extraction. Five or six independent transfection experiments were performed for each set of conditions.

### Ribosome profiling experiments

HeLa cells were plated on day 0 (D0) at a density of 1 million cells per plate in 10 mL MEM supplemented with 10% fetal bovine serum, 1% Gln, non-essential amino acids and a mixture of antibiotics and antimycotics (Gibco). On D+1, 80 μM TLN468 or 722 μM G418 was added. The cells were collected on D+2. The medium was removed, and the plates were placed on a liquid nitrogen bath, transferred to −70°C conditions before scratching. The cells were collected by adding polysome extraction buffer (10 mMTris CH_3_COONa pH7.6; 10 mM (CH_3_COO)_2_Mg; 10 mM NH_4_Cl; 1% Triton; 2 mM DTT).

Polysomes were extracted by adding 2x complete EDTA-free protease inhibitor (Roche) and 1 U per microliter murine RNase inhibitor (Biolabs ref: MO314S). Ribosome-protected fragments (RPF) were generated by 1 h of digestion with 15 U RNase I (Ambion)/OD_260nm_ at 25°C). Monosomes were separated by centrifugation on a 24% sucrose cushion at +4°C and were treated with DNase I. RNA was extracted with phenol at 65°C, CHCL_3_ and precipitated with 0.3 M CH_3_COONa pH5.2 in ethanol before loading onto a 17% acrylamide-bisacrylamide (19:1) gel containing 7 M urea and 1x Tris-acetate-EDTA (TAE). RPF at 28 to 34 nt were excised from the gel and precipitated in ethanol in the presence of glycogen.

RPF were depleted of ribosomal RNA with the Ribo-zero Human kit (Illumina) according to the manufacturer’s recommendations. The RPF libraries were constructed with the Biolabs NEBNext Multiplex Small RNA Library Prep Set for Illumina and sequenced on a NextSeq 500 High system with 75-base single reads.

### Synthesis of TLN468

#### 2,2,4-trimethyl-1,2-dihydroquinoline 1

An oven-dried Schlenk flask equipped with a nitrogen inlet and a magnetic stir bar was sequentially loaded with aniline (10 mmol), indium (III) chloride (0.5 mmol, 5 mol%) and acetone (12 mL). The mixture was stirred at 56°C until a full conversion of the starting material was observed. All the volatile components were removed under vacuum and the residue was purified by normal-phase column chromatography with a gradient of ethyl acetate in cyclohexane.

#### TLN468 (2)

To a solution of the corresponding 2,2,4-trimethyl-1,2-dihydroquinoline **1** (1 mmol) in dry acetonitrile (1 mL), we added 1 M HCl in diethyl ether (1 equivalent). Upon complete precipitation of the hydrochloride salt of **1**, the solids were collected by vacuum suction filtration on a fritted glass filter and repeatedly washed with hexanes. This material was used for the next step without further purification.

A round-bottomed flask equipped with a magnetic stir bar and a reflux condenser was loaded with the corresponding hydrochloride salt of **1** (1 mmol), dicyandiamide (1.01 equivalents), ethanol (1 mL) and water (1.5 mL). The mixture was refluxed for 18 h, then cooled to room temperature and the pH was adjusted to 9-10. The precipitate was collected by vacuum suction filtration on a fritted glass filter and washed thoroughly with cold water. It was then purified by reverse-phase column chromatography with a gradient of acetonitrile in water supplemented with 0.1% v/v formic acid, to yield quinazolyl-2-guanidine **2** as a formate salt (Figure 7). The analytical data obtained were consistent with published reports (^1^H and ^13^C NMR) (Deiana *et al*, 2020).

**Figure 7.**
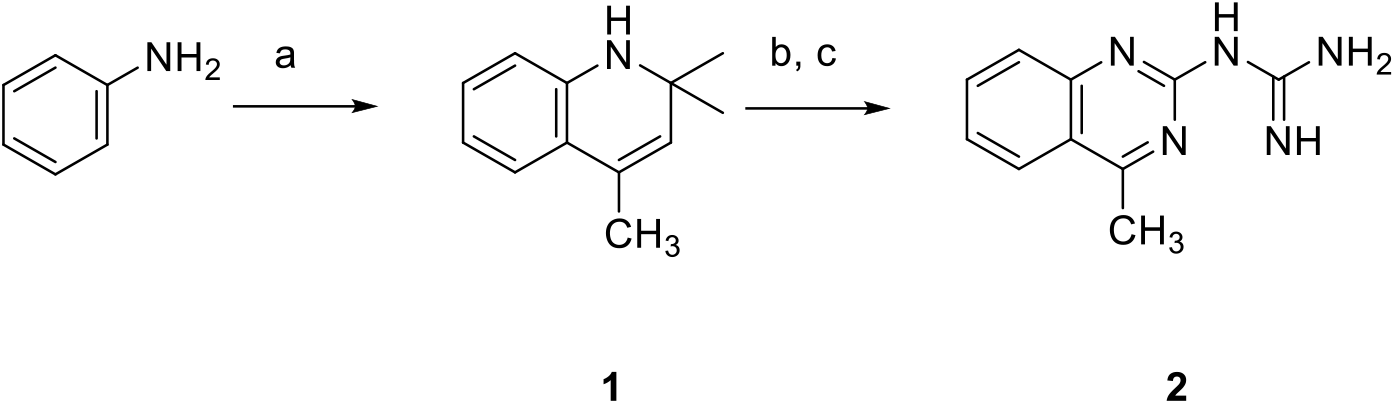
Chemical synthesis of TLN648. *Reagents and conditions*: (a) InCl_3_ (5 mol%), acetone, 56°C, 24-72 h; (b) 1 M HCl in diethyl ether (1 equivalent), acetonitrile, room temperature, 5-10 min; (c) dicyandiamide (1.01 equivalents), ethanol, H_2_O, reflux, 18 h.

## Acknowledgments

We thank the high-throughput sequencing facility of I2BC for library preparation and sequencing expertise. This work was funded by grants awarded to ON by the French foundation ARC (PJA20131200234), Vaincre la Mucoviscidose ON (No. RF20180502275), the ANR (grants: Rescue Ribosome (17-CE12-0024) Actimeth (19-CE12-0004-02)) and AFM-Telethon (grant 19660 and translectin No. 20531). OB was supported by the French foundation FRM (FDT20150532470). MC was supported by AFM-Telethon through a grant awarded to ON (grant 19660). EC was supported by AFM-Telethon through a grant awarded to JCC.

## Author contributions

OB and LB set up the reporter cell lines and all cell culture assays. OB, GM, and JCC performed HTS and analyzed the results. MC quantified readthrough efficiency for the PTCs of DMD. EC synthesized molecules for secondary assays. SK purified GST. DC performed mass spectrometry analysis. IH performed ribosome profiling experiments. PF performed the bioinformatics analysis. ON coordinated the work and analyzed the results. All authors participated in the writing of the manuscript.

## GEO databases

All RiboSeq data are available from the GEO database under accession number GSE185985.

---

**Table S1.** Frequency of the 40 most frequent DMD nonsense mutations

| Name | Nonsense mutation |  |  |  |  |  |  | Frequency (%) |
| --- | --- | --- | --- | --- | --- | --- | --- | --- |
| R145X | AGC | TGG | GTC | **TGA** | CAA | TCA | ACT | 0.75 |
| S147X | GTC | CGA | CAA | **TGA** | ACT | CGT | AAT | 0.14 |
| Q194X | TCA | GCC | ACA | **TAA** | CGA | CTG | GAA | 2.63 |
| R195X | GCC | ACA | CAA | **TGA** | CTG | GAA | CAT | 0.61 |
| Q267X | GAA | CAT | TTT | **TAG** | TTA | CAT | CAT | 2.63 |
| R539X | TTG | GGA | GAT | **TGA** | TGG | GCA | AAC | 0.34 |
| Q555X | GTT | CTT | TTA | **TAA** | GAC | ATC | CTT | 0.07 |
| L654X | TGG | GAT | ATT | **TAA** | CAT | CAA | AAA | 0.07 |
| E761X | GAC | TTA | AAA | **TAA** | AAA | GTC | AAT | 0.55 |
| R768X | GCC | ATA | GAG | **TGA** | GAA | AAA | GCT | 0.82 |
| R1051X | AAT | AAA | CTC | **TGA** | AAA | ATT | CAG | 0.55 |
| W1075 | AAG | GAG | GAA | **TAG** | CCT | GCC | CTT | 2.63 |
| E1182X | GAG | TAT | CTT | **TAG** | AGA | GAT | TTT | 0.07 |
| W1268X | TGG | GCA | TGT | **TGA** | CAT | GAG | TTA | 0.07 |
| R1577X | CGT | AAG | ATG | **TGA** | AAG | GAA | ATG | 0.68 |
| R1666X | GTC | ACC | TCC | **TGA** | GCA | GAA | GAG | 0.75 |
| R1844X | GAG | AGA | AAG | **TGA** | GAG | GAA | ATA | 0.27 |
| R1868X | AGG | TCT | CAA | **TGA** | AGA | AAA | AAG | 0.61 |
| W1879X | TCT | CAT | CAG | **TGA** | TAT | CAG | TAC | 0.07 |
| Y1882X | TGG | TAT | CAG | **TAA** | AAG | AGG | CAG | 0.14 |
| W1956X | AGC | AAG | CGC | **TAG** | CGG | GAA | ATT | 0.07 |
| R1967X | GCT | CAG | TTT | **TGA** | AGA | CTC | AAC | 0.61 |
| E2035X | TTT | AAG | CAA | **TAG** | GAG | TCT | CTG | 5.26 |
| R2095X | TAC | AAG | GAC | **TGA** | CAA | GGG | CGA | 0.55 |
| E2286X | ATA | AGC | CCA | **TAA** | GAG | CAA | GAT | 2.63 |
| Q2526X | AGG | CGT | CCC | **TAG** | TTG | GAA | GAA | 0.07 |
| R2553X | ATT | ACG | GAT | **TGA** | ATT | GAA | AGA | 0.82 |
| Q2574X | CGG | AGG | CAA | **TAG** | TTG | AAT | GAA | 2.63 |
| K2791X | GAA | CTT | CGG | **TAA** | AAG | TCT | CTC | 2.63 |
| R2870X | GAG | ACT | GTA | **TGA** | ATA | TTT | CTG | 0.82 |
| R2905X | CGG | CTT | CTA | **TGA** | AAG | CAG | GCT | 0.34 |
| W2925X | TCC | GCT | GAC | **TGA** | CAG | AGA | AAA | 0.14 |
| R2982X | AAG | GCA | CTT | **TGA** | GAA | GAA | AAT | 0.55 |
| R3034X | GTC | GAG | GAC | **TGA** | GTC | GTC | CAG | 0.55 |
| S3127X | TTG | AGC | CTG | **TGA** | GCT | GCA | TGT | 2.63 |
| R3190X | GAT | ACG | GGA | **TGA** | ACA | GGG | AGG | 0.89 |
| R3345X | TTT | TCT | GGT | **TGA** | GTT | GCA | AAA | 0.48 |
| R3370X | GAA | GAT | GTT | **TGA** | GAC | TTT | GCC | 1.02 |
| R3381X | AAC | AAA | TTT | **TGA** | ACC | AAA | AGG | 1.57 |
| R3391X | AAG | CAT | CCC | **TGA** | ATG | GGC | TAC | 1.5 |

**Figure S1.**
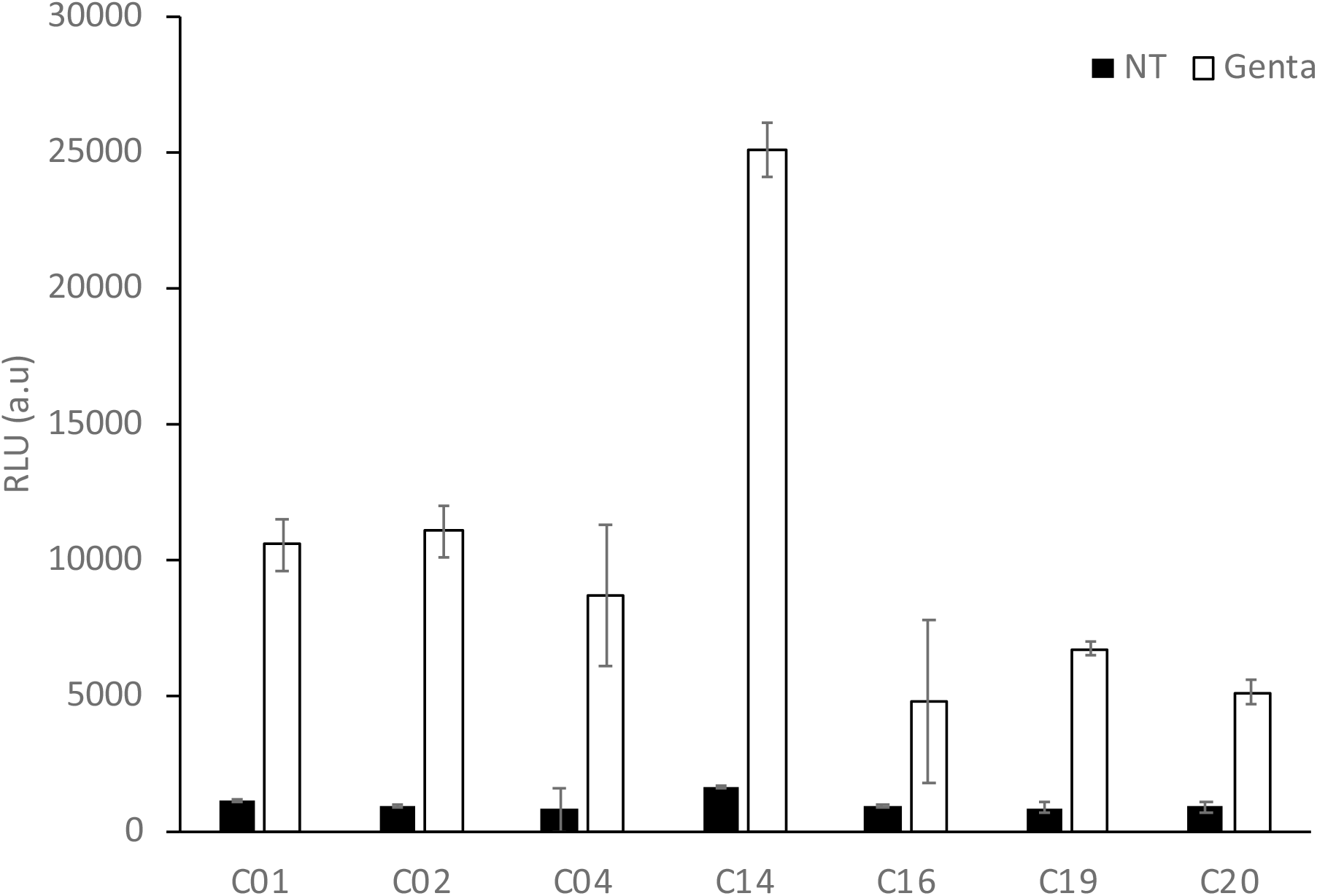
Selection of the recombinant cell line for HTS Recombinant NIH3T3 cells were cultured in 96-well plates, at an initial density of 15,000 cells/well. Luciferase activity was quantified by adding coelenterazine directly to the wells (*n*=4). Black and white bars represent basal *Metridia* luciferase activity and gentamicin (1.6 mM)-induced activity, respectively. We retained clone number C14 for further development.

